# RanGTP induces an effector gradient of XCTK2 and importin α/β for spindle microtubule cross-linking

**DOI:** 10.1101/664821

**Authors:** Stephanie C. Ems-McClung, Mackenzie Emch, Stephanie Zhang, Serena Mahnoor, Lesley N. Weaver, Claire E. Walczak

## Abstract

High RanGTP around chromatin is important for governing spindle assembly during meiosis and mitosis by releasing the inhibitory effects of importin α/β. Here we examine how the Ran gradient regulates Kinesin-14 function to control spindle organization. We show that *Xenopus* Kinesin-14, XCTK2, and importin α/β form an effector gradient, which is highest at the poles that diminishes toward the chromatin and is inverse of the RanGTP gradient. Importin α/β preferentially inhibit XCTK2 anti-parallel microtubule cross-linking and sliding by decreasing the microtubule affinity of the XCTK2 tail domain. This change in microtubule affinity enables RanGTP to target endogenous XCTK2 to the spindle. We propose that these combined actions of the Ran pathway are critical to promote Kinesin-14 parallel microtubule cross-linking at the spindle poles to cluster centrosomes in cancer cells. Furthermore, our work illustrates that RanGTP regulation in the spindle is not simply a switch, but rather generates effector gradients where RanGTP gradually tunes the activities of spindle assembly factors.

**Summary:** Ems-McClung et al. visualize a RanGTP effector gradient of association between XCTK2 and importin α/β in the spindle. The importins preferentially inhibit XCTK2-mediate anti-parallel microtubule cross-linking and sliding, which allows XCTK2 to cross-link parallel microtubules and help focus spindle poles.

## Introduction

The small GTP-binding protein, Ran, is required for microtubule (MT) nucleation and spindle assembly (Carazo-Salas et al., 1999; Kaláb et al., 1999; Ohba et al., 1999; Wilde and Zheng, 1999). RanGTP forms a gradient around chromatin that diminishes toward spindle poles (Kaláb et al., 2006; Kaláb et al., 2002) due to the localization of its guanine nucleotide exchange factor, RCC1, on chromatin (Moore et al., 2002). RanGTP acts through the nuclear transport receptors, importin α/β, which bind to NLS containing proteins and inhibit their activity in areas of low RanGTP. However, in areas of high RanGTP, this inhibition is released, resulting in localized activation of the NLS containing proteins that can stimulate microtubule (MT) nucleation and dynamics. It is postulated that the RanGTP gradient will result in the formation of downstream effector gradients of these MT regulators (Athale et al., 2008; Caudron et al., 2005), but there has been no direct visualization of these effector gradients. It is important to note that based on this model, RanGTP interaction with importin β would only happen near the chromatin resulting in the local release and activation of NLS containing proteins that dissipates toward the poles (Caudron et al., 2005). However, whether all NLS containing proteins will respond equivalently to the RanGTP gradient is unknown. In addition, the RanGTP gradient can induce feedback loops by the generation of increased numbers of microtubules, which are bound by components of the Ran pathway, resulting in localized effects on the spindle (Oh et al., 2016).

Key effectors of the RanGTP gradient include the nuclear transport receptors, importin α/β, as well as spindle assembly factors (SAFs) that are normally sequestered by importin α/β (Kaláb and Heald, 2008). These SAFs include a number of MT associated proteins, such as TPX2 (Gruss et al., 2001), NuMA (Nachury et al., 2001; Wiese et al., 2001), HURP (Koffa et al., 2006), and NuSAP (Ribbeck et al., 2006), as well as molecular motor proteins, Kid (Tahara et al., 2008; Trieselmann et al., 2003) and XCTK2 (Ems-McClung et al., 2004) that contain binding sites for either importin α and/or importin β. XCTK2 is a minus-end directed Kinesin-14 motor that cross-links and slides both parallel and anti-parallel MTs (Hentrich and Surrey, 2010) and contributes to proper spindle assembly, spindle length, and spindle pole formation (Cai et al., 2009; Walczak et al., 1997; Walczak et al., 1998). XCTK2 association within the spindle is spatially controlled by the RanGTP gradient through the tail domain (Hallen et al., 2008; Weaver et al., 2015), suggesting that the RanGTP gradient may specifically regulate the MT binding properties of XCTK2 to the spindle. How the RanGTP gradient regulates the localization of SAFs to the spindle is unknown.

Kinesin-14 proteins play critical roles in spindle pole formation in multiple organisms (Fink et al., 2009; Goshima et al., 2005a; Goshima et al., 2005b; Hatsumi and Endow, 1992b; Matuliene et al., 1999; Mountain et al., 1999; Sharp et al., 1999; She and Yang, 2017) and may be especially important in cells with centrosome amplification (Kwon et al., 2008), where they use their minus-end directed motor activity to help cluster those centrosomes into a bipolar spindle. Indeed, small molecule inhibitors that target the human Kinesin-14, HSET, may be useful as targeted therapeutics in cancers with centrosome amplification (Watts et al., 2013); therefore, understanding their mechanism of action in the spindle is an important avenue of pursuit. One conundrum is to understand how a molecular motor that cross-links and slides both anti-parallel and parallel MTs through its motor and tail domains can be involved in MT focusing at the spindle poles, where the NLS site in the tail would be predicted to be bound by importin α/β. Here, we develop XCTK2 biosensors that visualize a gradient of association with importin α in the spindle that is highest at spindle poles. We show that the importins preferentially inhibit XCTK2-mediated anti-parallel MT cross-linking and sliding and propose that importin α/β association with XCTK2 near the spindle poles is an effector gradient, inverse of the Ran gradient, that functions to facilitate parallel MT cross-linking and sliding for pole focusing. This model provides a mechanism by which XCTK2 localization and activity in the spindle is spatially controlled.

## Results and Discussion

The RanGTP gradient is proposed to impact the ability of importin α/β to interact with SAFs around the chromatin where RanGTP levels are high (Kaláb et al., 2006; Kaláb et al., 2002). We hypothesized that this would create an inverse effector gradient of importin α/β association with XCTK2. To test this idea, we developed FRET biosensors as a read out of the Ran-regulated interaction of importin α/β with XCTK2 in which we tagged importin α with CyPet (Imp α-CyPet) and full-length XCTK2 with YPet (YXCTK2) (Fig. 1). YXCTK2 produced a strong FRET signal at 525 nm with importin α-CyPet and importin β, indicating their association (Fig. 1 A). This association was disrupted by the addition of the RanGTP analog, RanQ69L (Fig. 1 B). In contrast, mutation of the NLS in XCTK2 (YXNLS) did not display FRET with importin α-CyPet and importin β, demonstrating their lack of association (Fig. 1 C). These results demonstrate that we have developed FRET probes that monitor the effects of RanGTP on XCTK2 and importin α/β association that can be used to look at effector gradients in the spindle.

**Figure 1.**
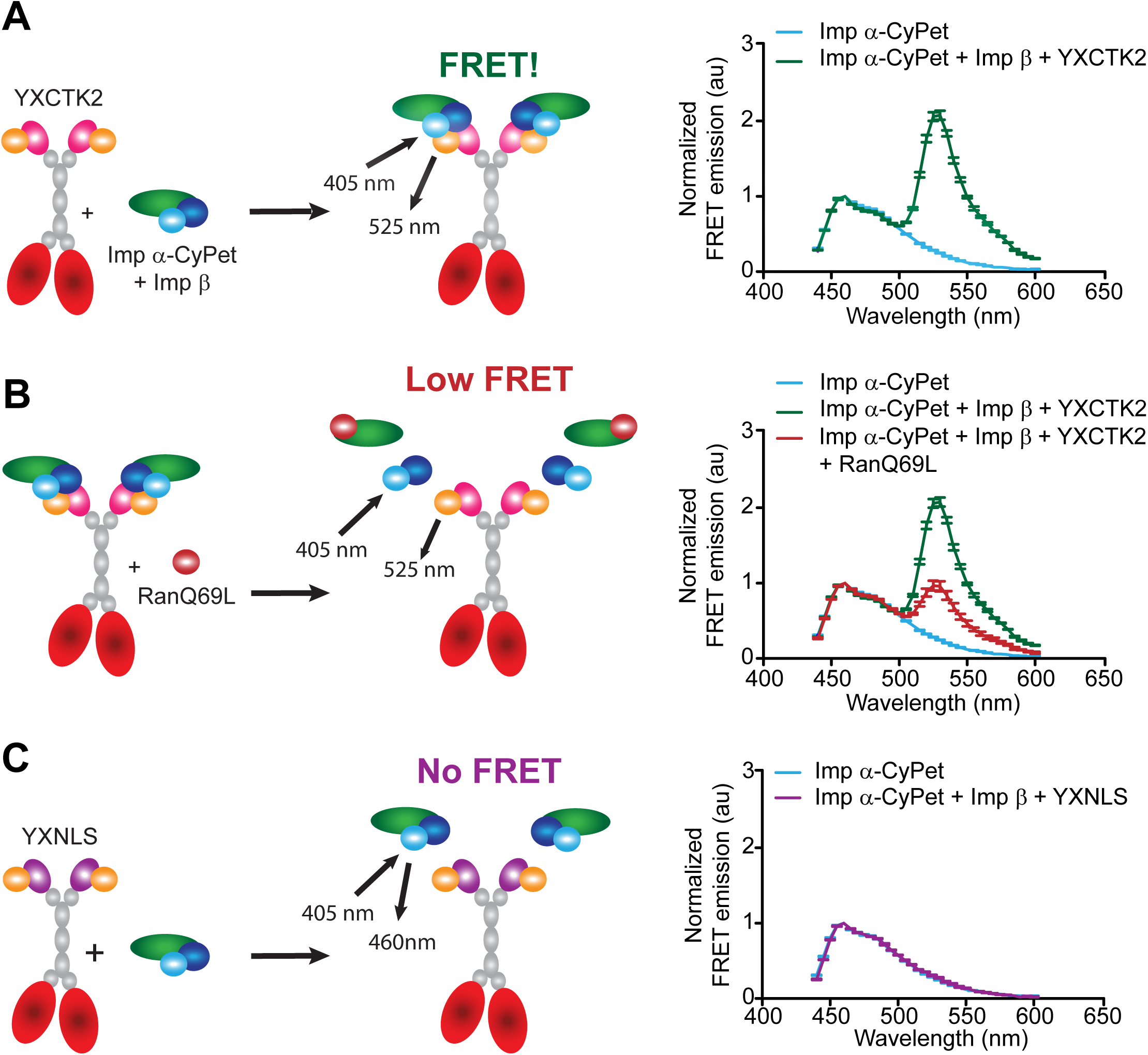
FRET biosensors recapitulate Ran-regulated association of XCTK2 and importin α/β. **(A-C)** Schematic (left) and solution-based FRET assay (right) with YXCTK2 and importin α-CyPet/importin β ± RanQ69L **(A**, **B)** or YXNLS and importin α-CyPet/importin β **(C)**. The normalized FRET ratios are graphed as mean ± SEM from 440-600 nm (*n* = 3 independent experiments).

To test whether there is a gradient of importin α/β association with XCTK2 within the spindle, we added importin α-CyPet with and without YPet tagged YXCTK2 or YXNLS to spindle assembly reactions in *Xenopus* egg extracts and imaged for CyPet, YPet, and rhodamine-MT fluorescence followed by fluorescence lifetime imaging microscopy (FLIM) (Fig. 2 A). Importin α-CyPet was generally diffuse in the cytoplasm with slight accumulation on the spindle; whereas, both YXNLS and YXCTK2 concentrated on the spindle to similar extents with localization enriched toward the poles, similar to the endogenous localization of XCTK2 (Fig. 2, A and B) (Ems-McClung et al., 2004; Walczak et al., 1997). The FLIM images of importin α-CyPet with YXCTK2 appeared to have shorter lifetimes than those of importin α-CyPet with YXNLS (Fig. 2 A), suggesting that there is FRET between importin α-CyPet with YXCTK2 on the spindle. We then used line scans to look at the distribution of lifetimes across the spindle and found that importin α-CyPet and importin α-CyPet with YXNLS had relatively flat distributions with mean lifetimes of 1.67 ± 0.01 ns and 1.68 ± 0.01 ns, respectively (Fig. 2 C). This result is consistent with our solution-based FRET assay in which YXNLS does not display FRET with importin α-CyPet and demonstrates that there is no interaction of YXNLS and importin α-CyPet on the spindle. In contrast, importin α-CyPet with YXCTK2 showed shorter lifetimes within the spindle demonstrated by a dramatic decrease in the average lifetime line scan across the spindle, 1.55 ± 0.01 ns (Fig. 2, A and C), indicating that XCTK2 interacts with importin α-CyPet within the spindle. In addition, the lifetimes for importin α-CyPet with YXCTK2 were shorter towards the spindle poles (Fig. 2, A and C) and coincided with the peak localization of YXCTK2 at the spindle poles, suggesting a gradient of association of importin α with XCTK2 extending from the poles to the chromatin (Fig. 2 D). The difference between the chromatin and pole lifetimes for YXCTK2 was statistically greater than either importin α-CyPet alone or importin α-CyPet with YXNLS, consistent with the idea of a gradient of association of importin α with YXCTK2, (Fig. 2 E; and Table S1). These results strongly suggest that XCTK2 and importin α form a gradient of interaction from the poles to the chromatin that is inverse from the RanGTP gradient.

**Figure 2.**
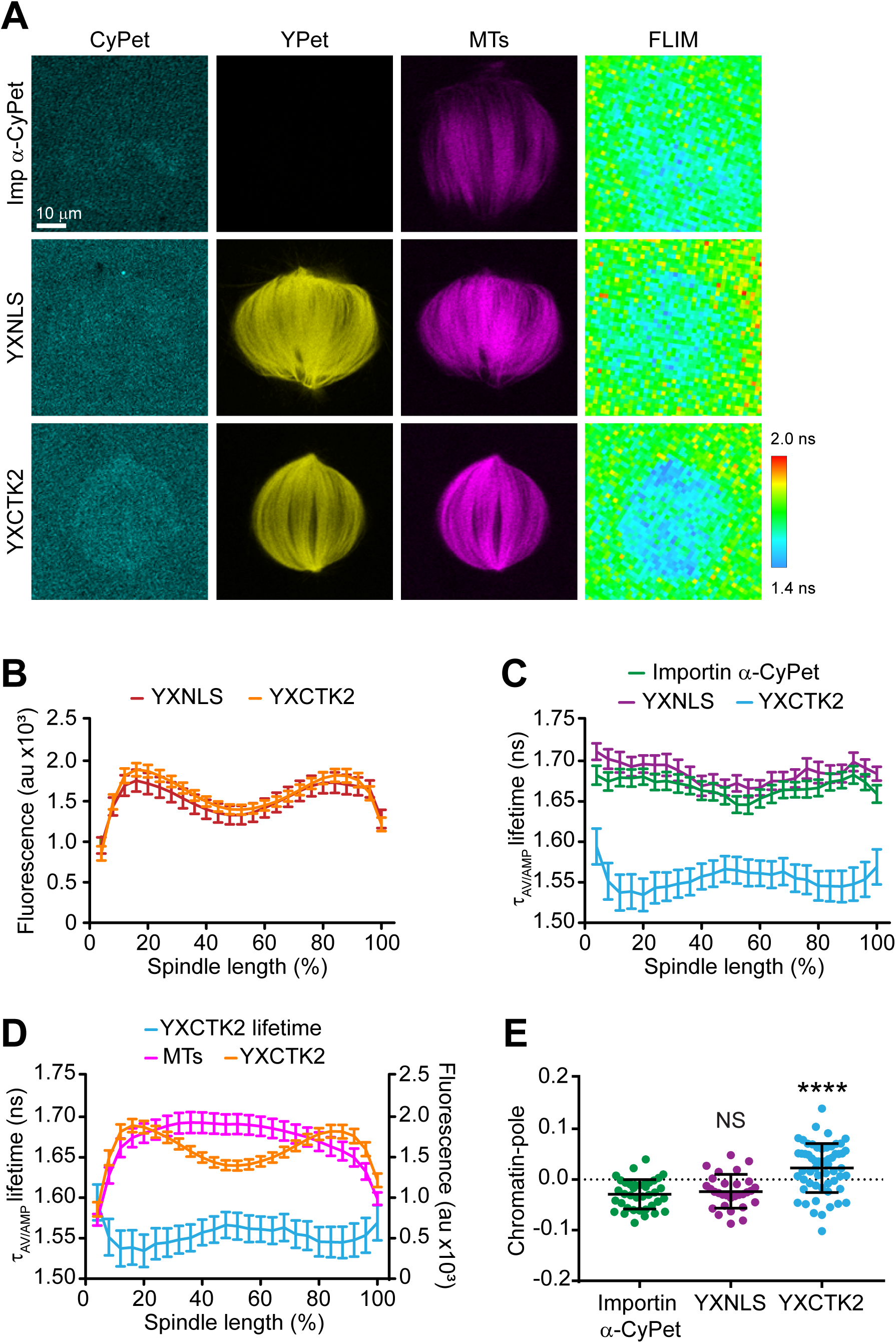
XCTK2 forms a gradient of association with importin α/β from the poles to the chromatin. **(A)** Representative confocal fluorescence (CyPet, YPet, and MTs) and lifetime (FLIM) images of importin α-CyPet + non-tagged XCTK2, importin α-CyPet + YXNLS, and importin α-CyPet + YXCTK2 spindle assembly reactions. Scale bar: 10 μm. **(B)** YPet fluorescence line scans of spindles assembled in **A** of importin α-CyPet with YXNLS or YXCTK2 normalized to percent spindle length (25 bins). YPet fluorescence percent spindle length is graphed as the mean ± SEM (*n* = 30 YXNLS + importin α-CyPet and 58 YXCTK2 + importin α-CyPet spindles from 3 independent experiments). **(C)** Lifetime line scans of spindles imaged in **A** and **B** where lifetimes are represented as the amplitude averaged lifetime (τ_AV/AMP_), and lifetimes per normalized percent spindle length are graphed as the mean ± SEM (*n* = 38 importin α-CyPet + XCTK2 spindles, 30 YXNLS + importin α-CyPet, and 58 YXCTK2 + importin α-CyPet spindles). **(D)** Line scans of the lifetimes from spindles with YXCTK2 + importin α-CyPet relative to the fluorescence of the spindle MTs and the YPet fluorescence plotted as in **B** and **C**. **(E)** Chromatin and pole lifetime differences for each spindle analyzed in **C** with the mean ± SD indicated (one-way ANOVA with Tukey’s multiple comparisons test: NS, not significant, **** *P* < 0.0001).

One way that RanGTP could modulate localization of SAFs within the spindle is by modulating the affinity of interactions of SAFs with importin α/β or with spindle MTs. We hypothesized that importin α/β binding could fine-tune XCTK2 function within spindles through the Ran-regulated association of the importins to the nuclear localization signal (NLS) in the XCTK2 tail domain. To test this idea, we designed additional FRET biosensors that monitored importin α/β binding to XCTK2 (Fig. S1, A and B) and used FRET as a readout to measure the binding affinity of importin α-YPet (Imp α-YPet) to CyPet tagged XCTK2 (CXCTK2) in the presence and absence of importin β (Fig. S1 C). Importin β significantly increased the affinity of importin α by decreasing the *K*_d_ five-fold and increasing the total amount bound (B_max_) by 28% (Fig. S1, C and D), consistent with previous studies showing that importin β releases the auto-inhibition of importin α for binding to NLS containing proteins (Catimel et al., 2001). To test how importin α/β binding to the XCTK2 tail modulates its interaction with MTs, we generated a construct containing only the XCTK2 tail domain tagged with YPet (YXTail), and used it to measure the affinity of tail MT binding in the presence and absence of an equal (1x) or four-fold molar excess (4×) of importin α/β (Fig. S1, E and F). Interestingly, addition of an equal molar amount of importin α/β did not change the *K*_d_, but decreased the amount bound by 46% (Fig. S1 F). In contrast, a four-fold excess of the importins increased the *K*_d_ by 10-fold and reduced the amount bound by 53% (Fig. S1 F). These results demonstrate that the XCTK2 tail MT affinity is tunable by importin α/β and provides a mechanism by which RanGTP could control the localization of XCTK2 within the spindle.

Kinesin-14s cross-link and slide both parallel and anti-parallel MTs (Braun et al., 2017; Fink et al., 2009; Hentrich and Surrey, 2010), but how importin α/β regulate these activities is unknown. We devised an *in vitro* assay using polarity marked MTs to assess how importin α/β modulate XCTK2 MT cross-linking and sliding (Fig. 3, A and B; Supplementary Videos 1, 2). Addition of a molar excess of importin α/β inhibited overall XCTK2 MT cross-linking, sliding, and pivoting (Fig. 3, C and D; and Table S2), consistent with inhibition of tail MT binding by importin α/β shown above. XCTK2 cross-linked both anti-parallel and parallel MTs to similar extents (*P* = 0.36) (Fig. 3 E; and Table S2), consistent with previous studies (Fink et al., 2009; Hentrich and Surrey, 2010). Surprisingly, importin α/β preferentially inhibited anti-parallel MT cross-linking by 45% with only a modest effect on parallel MT cross-links (Fig. 3 E; Table S2). The most dramatic effect of importin α/β was on the cross-linked MT populations that were actively sliding (Fig. 3 D). In this context, the effects of importin α/β on MT sliding were much more pronounced on anti-parallel MT cross-links (Fig. 3 F; and Table S2) relative to parallel MT cross-links (Fig. 3 G; and Table S2). These results are interesting because they suggest that the tail of XCTK2 has two different modalities of binding MTs and that importin α/β preferentially modulates the anti-parallel MT cross-linking and sliding activity.

**Figure 3.**
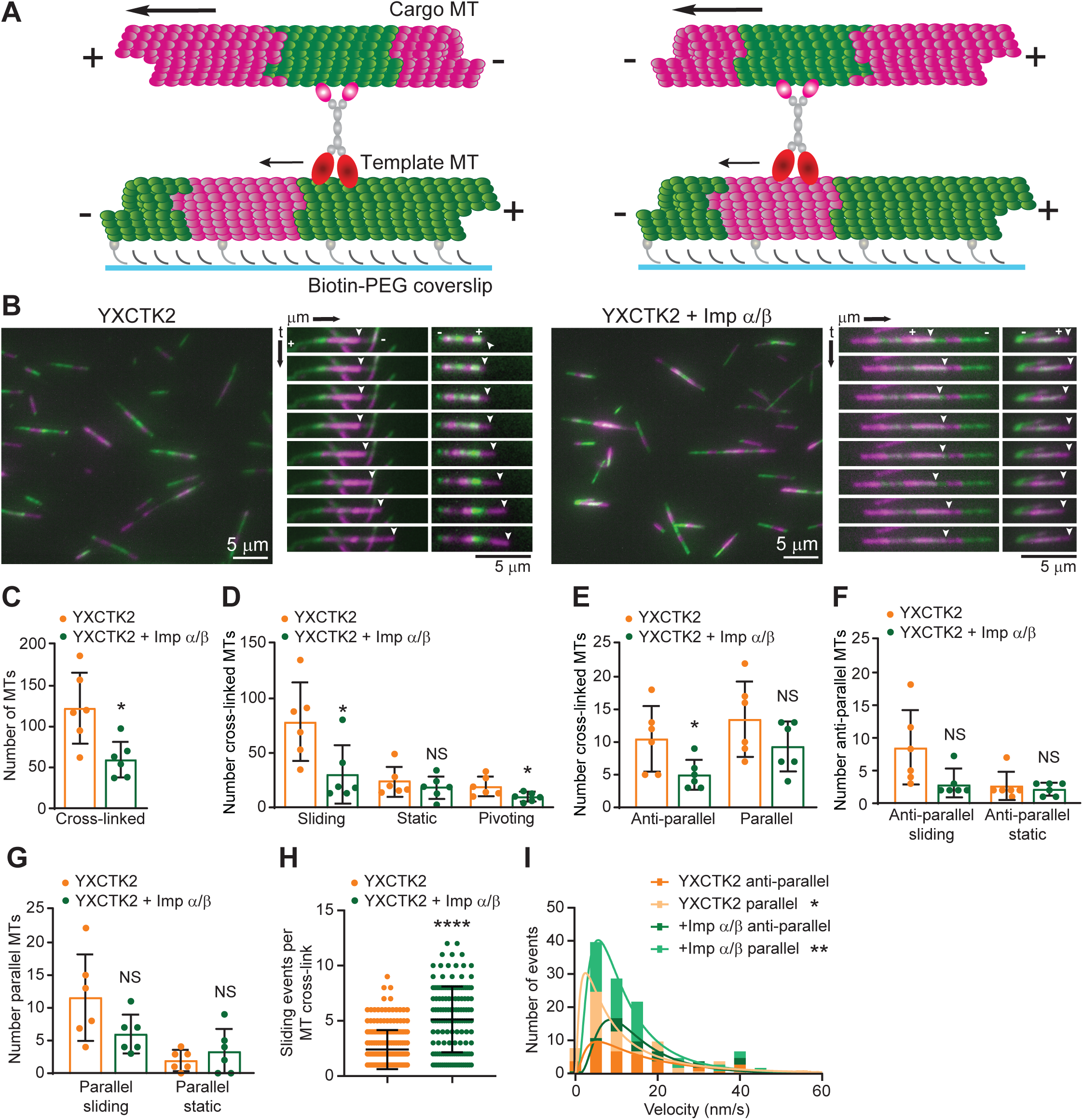
Importin α/β inhibit XCTK2 anti-parallel MT cross-linking and sliding. **(A)** Schematic of XCTK2 anti-parallel (left) and parallel (right) MT cross-linking and sliding assay with segmented MTs. **(B)** Representative images and kymographs of MT cross-linking and sliding with YXCTK2 (left) or YXCTK2 with 4-8 molar excess of importin α/β (right). The template MT minus (-) and plus (+) ends are indicated on the top kymograph image, and the sliding cargo MT plus end is indicated by an arrowhead. The YXCTK2 kymograph images increment by 30 sec intervals, and the YXCTK2 + importin α/β increment by 75 sec intervals to illustrate the multiple events over the course of the time-lapse. Scale bars: 5 μm. **(C-G)** Quantification of the indicated conditions plotted as the number of cross-links per experiment with the mean ± SD indicated as a bar graph with the dots representing the values of individual experiments (*n* = 729 YXCTK2 and 352 YXCTK2 + importin α/β cross-linked MTs, *n* = 63 YXCTK2 and 31 YXCTK2 + importin α/β anti-parallel cross-links, *n* = 81 YXCTK2 and 56 YXCTK2 + importin α/β parallel cross-links from 6 independent experiments; two-tailed unpaired Student’s *t*-test: NS, not significant, \**P* < 0.05). **(H)** The number of sliding events per MT cross-link are graphed with the mean ± SD (*n* = 212 YXCTK2 and 140 YXCTK2 + importin α/β cross-links from 3 independent experiments; Welch’s two-tailed *t*-test: **** *P* < 0.0001). **(I)** Velocity of MT sliding events for anti-parallel and parallel MT cross-links are graphed as frequency histograms with the best fit log Gaussian curves (*n* = 42 anti-parallel events and71 YXCTK2 parallel, and *n* = 60 YXCTK2 + importin α/β anti-parallel and 116 parallel events from 3 independent experiments; extra sum-of-squares F-test to compare geometric means of the log Gaussian distribution of events: * *P* < 0.05, ** *P* < 0.01; YXCTK2 parallel vs. YXCTK2 + importin α/β parallel *P* = 0.4299; YXCTK2 anti-parallel vs. YXCTK2 + importin α/β anti-parallel *P* = 0.4020). The frequency histogram is graphed up to 60 nm/s, excluding a single point at 120 nm/s.

Cargo MTs often slid stochastically along the template MTs, being interrupted by pausing, reversing directions and/or changing their rate of sliding. Within the 10 min time frame of the movies, the cargo MTs on average had two distinct sliding events per MT cross-link (Fig. 3 H; Video 1; and Table S2). Addition of importin α/β increased the number of sliding events per MT cross-link two-fold (Fig. 3 H; Video 2; and Table S2). The increased sliding events in the presence of importin α/β are consistent with reduced tail MT affinity that could result in more frequent release of the tail domain from MTs due to competition with importin α/β. XCTK2 also slid anti-parallel MTs 41% faster than parallel MT cross-links (Fig. 3 I; Video 1; and Table S2), consistent with differences in sliding observed previously (Hentrich and Surrey, 2010). Importin α/β addition did not affect the sliding velocity of either population of sliding events (Fig. 3 I; Video 2; and Table S2), suggesting that the tail domain cannot bind MTs and importin α/β simultaneously, which implies that importin α/β simply compete with MTs for binding to the tail domain (Chang et al., 2017).

Previous work showed that the tail domains of Kinesin-14 motors are important for spindle localization (Hallen et al., 2008; Weaver et al., 2015), and thus differential interaction with importin α/β may affect XCTK2 localization. To look at the effects of RanGTP gradient on XCTK2 localization within the spindle, we enhanced the RanGTP gradient in *Xenopus* egg extracts (Halpin et al., 2011) and quantified the localization of endogenous XCTK2. Adding purified Ran to spindle assembly reactions enhanced the RanGTP gradient, resulting in steeper gradients with higher RanGTP around the chromatin that decreased to control levels at the spindle poles as visualized by Rango-2, a RanGTP gradient biosensor (Fig. 4, A-C; and Table S3). Enhancing the RanGTP gradient by the addition of 10 μM or 20 μM Ran increased the overall localization of endogenous XCTK2 to the spindle by 12% and 17%, respectively (Fig. 4, D and E; and Table S4). Enhancing the RanGTP gradient did not change the overall morphology of the spindles, as the MT polymer level, spindle area, eccentricity, length, and width were largely unchanged, as well as the distribution of MT polymer across the spindle (Fig. S2,A-F; and Table S4). These results suggest that the RanGTP gradient promotes the global localization of Kinesin-14s to the spindle.

**Figure 4.**
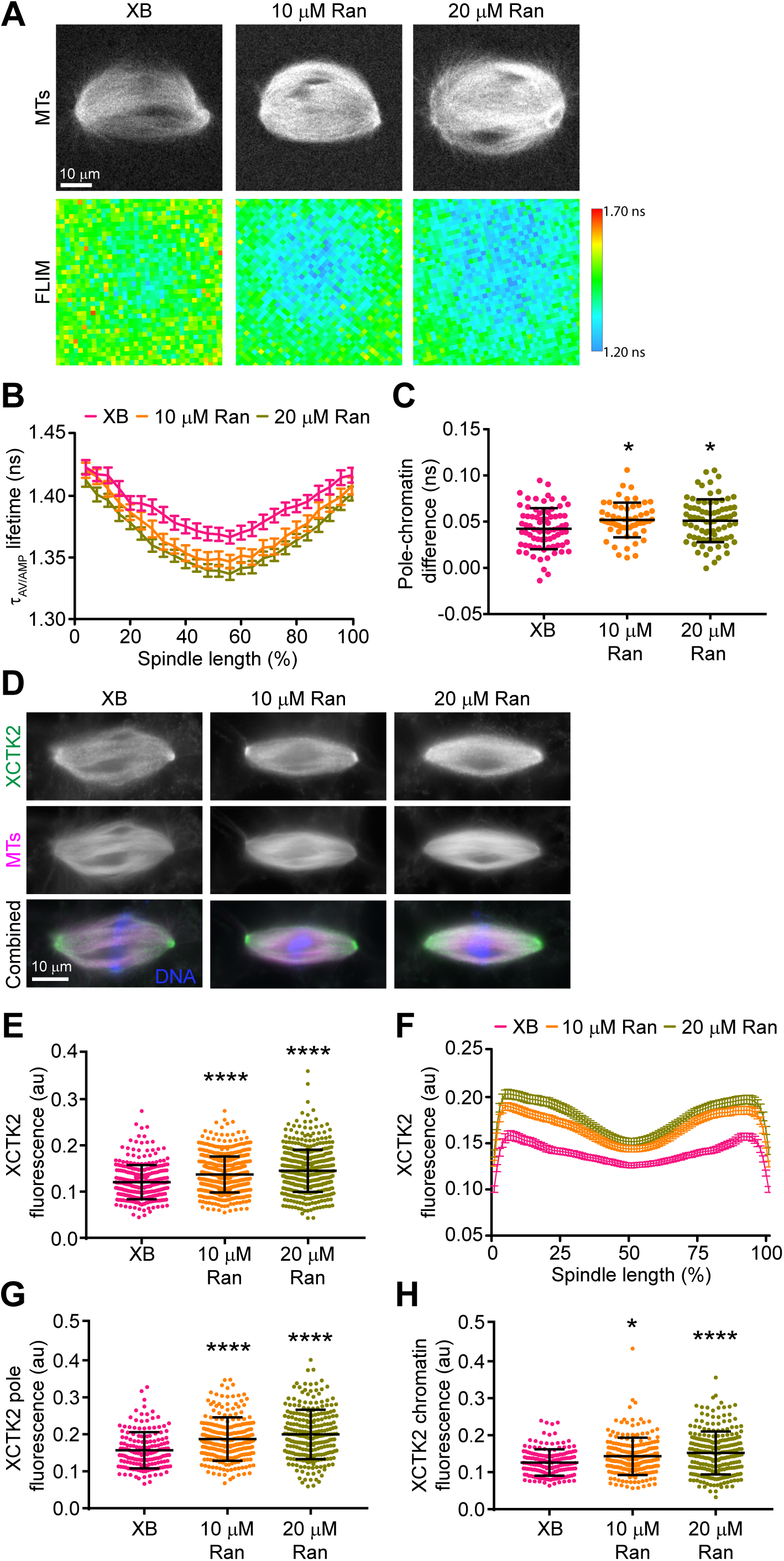
The RanGTP gradient promotes global and local targeting of XCTK2 within the spindle. **(A)** Representative MT confocal and Rango-2 FLIM images of spindles assembled with XB control buffer, 10 μM Ran, or 20 μM Ran addition. Lifetimes are represented as the amplitude averaged lifetime (τ_AV/AMP_). **(B)** Line scans from pole to pole of the spindle lifetime images for the indicated conditions normalized for percent spindle length (25 bins). Lifetimes per spindle length are graphed as the mean ± SEM (*n* = 57-78 spindles per condition from 3-4 independent experiments). **(C)** Lifetime difference between the pole and chromatin regions for each spindle in each condition from **B** (one-way ANOVA with Tukey’s multiple comparisons test: * *P* < 0.05). **(D)** Representative wide-field fluorescence microscopy images of spindle assembly reactions from parallel experiments with XB control buffer or with added Ran that were stained with α-XCTK2. **(E)** Spindles from **D** were analyzed using a Cell Profiler pipeline that measured the mean total XCTK2 spindle fluorescence based on α-XCTK2 staining in the indicated conditions and plotted with the mean ± SD (*n* = 314-564 spindles per condition from 5 independent experiments). **(F)** Line scans of bipolar spindles from **D** were performed, normalized for percent spindle length (101 bins), and graphed as the mean ± SEM (*n* = 168-248 spindles per condition from 4 independent experiments). **(G)** XCTK2 peak pole fluorescence plotted as the average fluorescence from the two poles per spindle analyzed in **F**. **(H)** XCTK2 chromatin fluorescence plotted as the center spindle position for each spindle analyzed in **F**. In **G** and **H**, the mean ± SD is indicated. **(E-H)** Kruskal-Wallis test with Dunnett’s Multiple Comparisons test compared to XB control, * *P* < 0.05, **** *P* < 0.0001. Scale bars: 10 μm.

Kinesin-14 localization is enriched toward the spindle poles where it functions in bipolar spindle assembly (Hatsumi and Endow, 1992a; Walczak et al., 1997). Enhancing the RanGTP gradient showed an overall increase in XCTK2 localization across the spindle that was significantly enriched toward the spindle poles relative to the chromatin region, suggesting that the RanGTP gradient also influences the localization of XCTK2 within the spindle (Fig. 4, F-H; and Table S4). Enhancing the RanGTP gradient resulted in a small but less dramatic change in the localization of NuMA, another Ran-regulated SAF, to the spindle (Fig. S2, G-K; and Table S4). These results suggest that the RanGTP gradient differentially controls the spatial localization of SAFs within the spindle. Setting up different effector gradients in the spindle would provide a mechanism for cells to respond to local changes in RanGTP levels.

Overall our results uncover several important findings about how the RanGTP gradient modulates SAF activity in the spindle. First, we found that a key action of Ran and the importins is to modulate the binding affinity of the XCTK2 tail to MTs, which targets XCTK2 to the spindle and regulates its spatial distribution (Fig. 5 A). This is consistent with our previous work, showing that XCTK2 turnover in the spindle was spatially controlled by Ran (Weaver et al., 2015). More intriguing is the current observation that the importins preferentially inhibit XCTK2 anti-parallel MT cross-linking and sliding (Fig. 5 A). Kinesin-14s can cross-link both parallel and anti-parallel MTs, but how a single MT binding domain can bind to a structural polymer in two orientations is not known. Studies with Drosophila Ncd have suggested that there are two regions of MT binding in the tail domain (Karabay and Walker, 1999; Karabay and Walker, 2003; Wendt et al., 2003). One interesting idea is that Kinesin-14s have two separate MT binding domains in the tail, one that mediates parallel MT cross-links and one that mediates anti-parallel MT cross-links. In this model, importin α/β would only inhibit the anti-parallel MT binding, leaving parallel MT cross-linking intact. This model would help explain how Kinesin-14s can focus MT minus ends at the spindle poles, which is an area of high importin α/β association (Fig. 5, A and B). In addition, this model is consistent with our biochemical analysis of XCTK2 tail MT binding, which is reduced but not abolished by the presence of importin α/β. Future structural studies will thus be critical to determine these different modes of MT binding.

**Figure 5.**
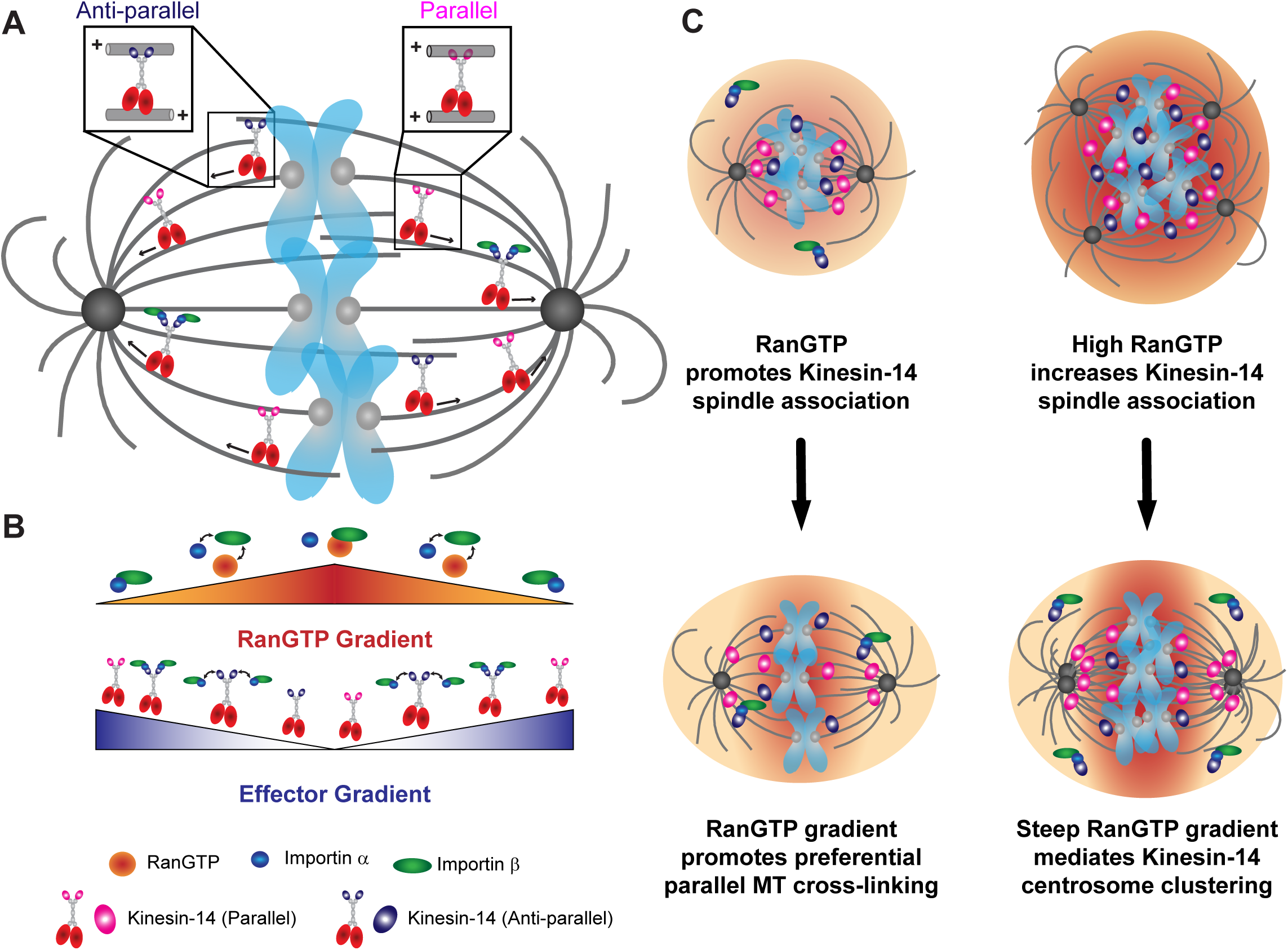
The RanGTP gradient generates an effector gradient of Kinesin-14s for preferential parallel MT cross-linking and sliding near the spindle poles. **(A)** In low amounts of importin α/β or high RanGTP near the chromatin, XCTK2 cross-links and slides both anti-parallel and parallel MTs. In high levels of importin α/β or low levels of RanGTP near the spindle poles, importin α/β selectively inhibit XCTK2 anti-parallel MT cross-linking and reduce the number of MTs that slide. **(B**, **C)** The RanGTP gradient sets up an inverse effector gradient of importin α/β association with XCTK2 that promotes pole focusing through preferential parallel MT cross-linking and sliding. In cancer cells with centrosome amplification and high RanGTP, we propose that RanGTP increases Kinesin-14 association to the spindle and that the RanGTP gradient sets up an effector gradient of Kinesin-14 localization and activity that mediates centrosome clustering for cancer cell survival.

RanGTP modulates several SAFs through the action of importin α/β, suggesting a common mechanism. In contrast, our work demonstrates that not all SAFs respond equivalently to the action of RanGTP in the spindle. Most notably, high RanGTP promoted both XCTK2 association with the spindle and an increase in localization toward the poles, but high RanGTP had only minor effects on NuMA localization, which is another Ran and importin α/β regulated SAF (Chang et al., 2017; Nachury et al., 2001; Wiese et al., 2001). The prevailing model on how the RanGTP gradient regulates SAFs is to release the inhibitory effects of importin α/β around chromatin, essentially setting up an effector gradient parallel to the RanGTP gradient. Our combined FLIM and cross-linking results support the idea that the RanGTP gradient sets up an inverse effector gradient of importin α/β and XCTK2 association within the spindle for preferential parallel MT cross-linking and sliding at the spindle poles rather than turning off XCTK2 activity near the poles (Fig. 5 B). One idea is that the RanGTP gradient in conjunction with importin α/β act as a rheostat to modulate several Ran-regulated effector gradients that will have altered activities rather than differential levels of activities as previously proposed. Furthermore, it is known that the Ran gradient is quite steep in the spindle (Athale et al., 2008), but perhaps the local interactions of importin α/β with the SAFs are more important for modulating activity, and thus would respond differentially to the steepness of the RanGTP gradient. In support of this idea, several components of the Ran pathway that influence MT behavior were shown to be controlled by their interaction with MTs, setting up a positive feedback loop that contributes to spindle assembly (Oh et al., 2016). Understanding differences in effector gradients and determining whether modulation of MT binding is a universal control mechanism by Ran will be important avenues for future studies.

Finally, our work brings important new insights into how cancer cells co-opt the spindle machinery for their own survival (Fig. 5 C). It is well known that many cancers have centrosome amplification, and centrosome amplification can lead to chromosome instability through clustering of centrosomes in multipolar spindles to allow for bipolar divisions and cell survival (Ganem et al., 2009; Silkworth et al., 2012). The human Kinesin-14 HSET is a key factor that promotes centrosome clustering in these cells (Chavali et al., 2016; Kwon et al., 2008; Watts et al., 2013). Various cancers also have increased Ran expression and enhanced RanGTP gradients due to increased DNA content (Hasegawa et al., 2013; Sheng et al., 2018; Xia et al., 2008). Some cancers also have increased HSET expression, which often correlates with metastasis and poor prognosis (Fu et al., 2018; Liu et al., 2016; Pannu et al., 2015; Patel et al., 2018). We propose that the enhanced RanGTP gradients from higher DNA content along with the increased HSET expression in cancer cells promote HSET localization to the spindle with a bias to the spindle poles that facilitates centrosome clustering (Fig. 5 C). Thus, cancer cells have taken advantage of their enhanced RanGTP gradient for their own benefit and survival.

## Methods and methods

### Protein expression and purification

Plasmids for the expression of 6His-Rango-2 (pRSETA-Rango-2) (Kaláb and Soderholm, 2010; Kaláb et al., 2002) and 6His-Ran (pRSETB-Ran) (Wilde and Zheng, 1999) were induced in BL21(DE3)pLysS bacteria and purified using NiNTA agarose (Qiagen) as previously described (Ems-McClung et al., 2004) except that protein expression was induced at 16 °C for 24h. Proteins were dialyzed into XB dialysis buffer (10 mM HEPES, pH 7.2, 100 mM KCl, 25 mM NaCl, 50 mM sucrose, 0.1 mM EDTA, 0.1 mM EGTA). Rango-2 was further purified by gel filtration. Proteins were aliquoted and flash frozen in liquid nitrogen.

To express the tail domain of XCTK2 fused to YPet, a parental pRSETB-YPet plasmid was created by PCR amplification of the YPET DNA from pRSETA-Rango-2 using GFPBamHIF2 and YPetSacIR2 primers and inserted into the *BamH*I/*Sac*I sites of pRSETB. The cDNA for the tail domain of XCTK2, amino acids 2-120, was inserted into the *Sac*I/*Hind*III sites of pRSETB-YPet for the expression of 6His-YPet-XCTK2-Tail (YXTail). To express importin α fused to CyPet, a parental pRSETB-CyPet plasmid was created by PCR amplification of the CyPET DNA from pRSETA-Rango-2 using GFPBamHIF1 and GFP(TGA)HindIII primers and inserted into the *Nco*I/*Hind*III sites of pRSETB. The cDNA for importin α (*Xenopus* importin α1a) was amplified from pQE70-importin α (Ems-McClung et al., 2004) with BamHI-Importin α-F and Importin α-XhoI-R primers, digested with *BamHI* and *Xho*I, and cloned into the pRSETB-CyPet *BamH*I and *Xho*I sites to generate pRSETB-importin α-CyPet (importin α-CyPet). For 6His-importin α-YPet (importin α-YPet) protein expression, YPet DNA from pRSETB-YPet was digested with *BamH*I and *Hind*III and inserted into the 3’ *Bgl*II and *Hind*III sites of pRSETB-importin α-CyPet. 6His-YPet-XCTK2-Tail (YXTail), importin α-CyPet, importin α-YPet, importin α-6His (importin α), and 6His-S-importin β (importin β) were induced, purified using NiNTA agarose, dialyzed, and aliquoted as described above for 6His-Ran except YXTail was dialyzed into XB250 dialysis buffer (10 mM HEPES, pH 7.2, 250 mM KCl, 25 mM NaCl, 50 mM sucrose, 0.1 mM EDTA, 0.1 mM EGTA).

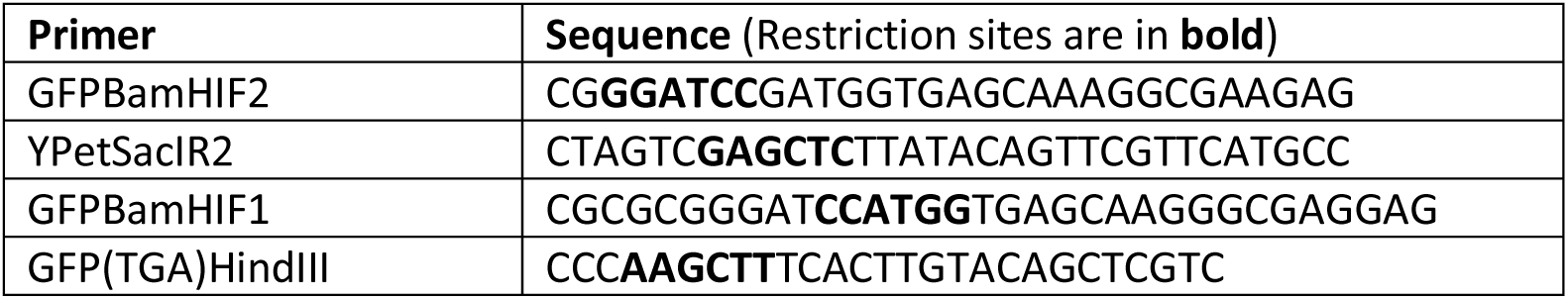
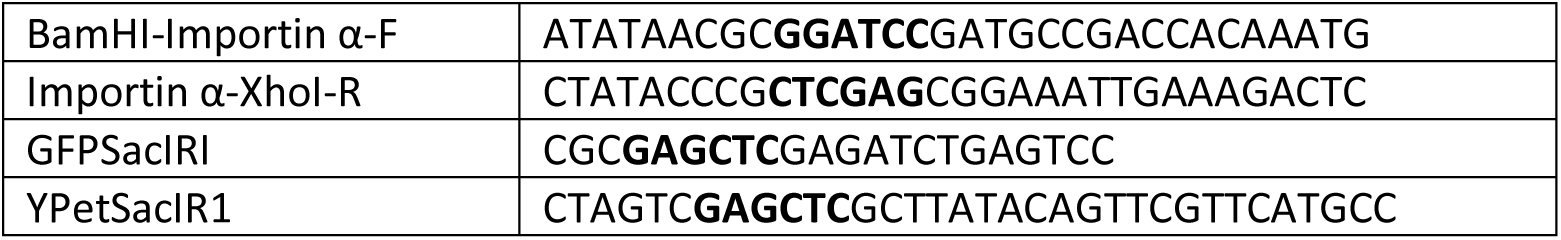

For expression of CyPet or YPet tagged full-length XCTK2, the DNA for CyPet or YPet was amplified with the GFPBamHIF1 and GFPSacIRI primers, digested with *BamH*I/*Sac*I, and inserted into pFastBac1-XCTK2 (Cai et al., 2009). For expression of YPet-XCTK2-NLS2 (YXNLS), YPet DNA was amplified using GFPBamHIF1 and YPetSacIR1 primers, digested with *BamH*I/*Sac*I, and inserted into pFastBac1-GFP-XCTK2-NLS2b (Cai et al., 2009), replacing the GFP DNA. Full-length XCTK2, CyPet-XCTK2 (CXCTK2), YPet-XCTK2 (YXCTK2), and YXNLS proteins were expressed using the Bac-to-Bac Baculoviral System (Invitrogen, Carlsbad, CA), and purified using traditional ion exchange and gel filtration chromatography as previously described (Ems-McClung et al., 2004). Final buffer composition was 20 mM PIPES pH6.8, 300 mM KCl, 1 mM MgCl_2_, 1 mM EGTA, 0.1 mM EDTA, 1 mM DTT, 10 uM MgATP, 0.1 μg/ml LPC, 10% sucrose. All proteins were flash frozen in small single use aliquots in liquid nitrogen, and quantified by absorbance using the extinction coefficient for CyPet (35,000 M^-1^ cm^-1^) or YPet (104,000 M^-1^ cm^-1^) and/or by densitometry of proteins electrophoresed on 10% SDS-PAGE gels using BSA as a control.

### Cycled spindle assembly and immunofluorescence

Cytostatic factor (CSF) extract was made from *Xenopus laevis* eggs as previously described (Murray, 1991) except that cytochalasin B (cytoB) was used in place of cytochalasin D. The CSF extract was supplemented with 1/1000 10 mg/ml LPC, 1/1000 cytoB; 1/50 50× energy mix (150 mM creatine phosphate, 20 mM ATP, 2 mM EGTA, 20 mM MgCl_2_), 1/40 2M sucrose, and 0.3 μM X-Rhodamine tubulin. CSF extracts containing 200 demembranated sperm/μl were cycled by the addition of 25x calcium chloride solution (final: 10 mM HEPES pH7.7, 1 mM MgCl_2_, 100 mM KCl, 150 sucrose, 10 μg/ml cytochalasin D, 10 mM CaCl_2_) at room temperature for approximately 60 min followed by the addition of an equal volume CSF extract.

For immunofluorescence, 25 μl of cycled extract was mixed with equal volumes of XB dialysis buffer or a 20x stock of Ran to give 10 μM or 20 μM final concentrations. Reactions were incubated at room temperature for 45 min before mixing with 30% glycerol, 0.5% Triton X-100, BRB80 (80 mM PIPES pH6.8, 1 mM MgCl_2_, 1 mM EGTA) and sedimenting onto coverslips over a 40% glycerol/BRB80 cushion. Coverslips were fixed with cold methanol for 5 min and rehydrated in TBS-Tx (10 mM Tris pH 7.6, 150 mM NaCl, 0.1% Triton X-100). Spindle assembly structures were immunostained by blocking with AbDil-Tx (2% BSA, TBS-Tx) and then incubated with 1 μg/mlα-XCTK2 (Walczak et al., 1997) or 2.5 μg/ml α-NuMA Tail II (Walczak et al., 1998) in AbDil-Tx for 30 min, washed with TBS-Tx, incubated with 1 μg/ml donkey anti-rabbit Alexa488 in AbDil-Tx for 30 min, washed with TBS-Tx, and then stained with 2 μg/ml Höechst in TBS-TX for 5 min. Coverslips were washed in TBS-Tx, mounted onto slides with Prolong Diamond (Invitrogen), and sealed with nail polish.

Immunofluorescent spindle assembly reactions were imaged on a Nikon A1 microscope mounted with a Hamamatsu camera and Plan Fluor 40x Oil DIC HN2 NA1.3 objective that was controlled by Elements (Nikon). Images for each condition were acquired in three different channels (DAPI, FITC or TRITC) as a 10×10 or 12×12 array using the Scan Large Image module. Each image was acquired with equivalent exposure times per channel. Images containing spindles were manually grouped per experiment per condition for analysis with a custom-built Cell Profiler pipeline. The MTs in the TRITC channel were enhanced for edges, smooth texture, and tubeness. Spindles were identified in the enhanced MT image by manually drawing a line inside the spindle. An outline was then created around the spindles using the IdentifySecondaryObjects module. The mean intensity of the TRITC (MTs) and FITC (XCTK2 or NuMA) channels were quantified, and the area, eccentricity, length, and width were calculated. Results were exported as an Excel file and graphed in Prism. Analysis of variance (ANOVA) tests were performed in Prism based on the normality of the samples.

The localization of XCTK2 or NuMA within the bipolar spindles identified in Cell Profiler was determined using line scans generated with the XLineScan plugin for Fiji (Wilbur and Heald, 2013). Briefly, 15-pixel wide lines were manually drawn from pole to pole of bipolar spindles on composite DAPI/FITC/TRITC images, and the FITC and TRITC channel fluorescence intensities were measured and normalized to 101 bins. The results were exported as an Excel file and analyzed in Prism 7. The differences between poles and chromatin fluorescence were calculated in Excel by taking the difference between the average of the three pole fluorescence values centered on the peak XCTK2 or NuMA fluorescence at both ends of the spindle (positions 6, 7, 8 and 94, 95, 96) from the average of the center three fluorescence values (positions 50, 51, 52) for each spindle for each condition and plotted in Prism. The peak fluorescence of XCTK2 and NuMA were used because the microtubule fluorescence was dome shaped and did not peak at the poles. Analysis of variance (ANOVA) tests were performed in Prism (GraphPad) based on the normality of the samples.

### Fluorescence lifetime imaging microscopy (FLIM)

Confocal and FLIM imaging of the Rango-2 RanGTP biosensor was performed in parallel cycled spindle assembly reactions except that sperm was added to the extract at a final concentration of 600 sperm μl^-1^. Reactions were supplemented with a 20x stock of proteins for a final concentration of 2 μM Rango-2 with XB dialysis buffer, 10 μM Ran, or 20 μM Ran and incubated for 45-60 min at room temperature (RT) before squashing 4 μl under a 22×22 mm #1.5 coverslip and imaging sequentially for confocal and FLIM images. A Leica SP8 confocal microscope equipped with a DMi8 inverted platform and a HC PL APO CS2 63x 1.2 NA water objective controlled by LAS-X software (Leica) was used to take confocal images of X-rhodamine fluorescence (MTs), and YPet and CyPet fluorescence (Rango-2). Images of MTs and Rango-2 were acquired with lasers at 40 MHz, as 256×256 pixel images at 400 MHz scanning with two-frame averaging, and a zoom factor of two. MTs and YPet were imaged with a white light laser at 70% power using the 594 nm laser line at 25% and the HyD5 PMT (610-700 nm) with variable gain for MTs and the 514 nm laser line at 50% and the SMD2 PMT (520-580 nm) with 50% gain for YPet. CyPet was imaged with a 440 nm laser at 90% power using SMD1 PMT (460-490 nm) with 500% gain. The pinhole was set to 1 AU or 111.5 μm.

FLIM of Rango-2 was performed sequentially after confocal imaging using an attached Pico Harp 300 (PicoQuant) time-correlated single photon counting system controlled by SymPhoTime64 software (PicoQuant) using a pulsed 440 nm laser at 20 MHz and 90% power. Images were acquired at 128×128 pixel at 200 MHz scanning and a zoom factor of two until the photon count reached 2000, which on average took 1.5 min. Spindle lifetimes were determined using the SymPhoTime64 software in which the decay data was binned 2×2 pixels and the overall decay of the image used to determine the model parameters since Rango-2 is soluble throughout the extract. Decay of CyPet in Rango-2 best fit a 3-exponential tail model based on the *X*^2^ values and the residuals. Lifetimes were then calculated per pixel by fixing the background and the individual lifetimes determined by the fitting. FLIM and photon count images were scaled identically in SymPhoTime and exported as bitmaps for color images, and TIFF or ASCII files for line scan analysis.

Spindle line scans were performed on color combined images in ImageJ using a XiaLineScan application (https://github.com/XiaoMutt/XiaoImageJApp/blob/master/store/XiaoImageJApp.jar) based on a similar application used by Wilbur and Heald (Wilbur and Heald, 2013). Briefly, for each FLIM acquisition in each condition and experiment, the confocal X-rhodamine, YPet, and CyPet images were scaled to 64×64 32-bit images and color combined with the corresponding FLIM image as a ZIP file. A 3-pixel width line was drawn from one spindle pole to the opposite pole based on the microtubule (X-rhodamine) channel and the average lifetimes from the FLIM channel recorded and normalized to a 25-pixel line length. Results were exported into Excel and graphed in Prism. The differences between poles and chromatin lifetimes were calculated in Excel by taking the difference between the average of the three pole lifetimes centered on the peak microtubule fluorescence at both ends of the spindle (positions 2, 3, 4 and 22, 23, 24) from the average of the center three lifetimes (positions 12, 13, 14) for each spindle for each condition and plotted in Prism. ANOVA tests were performed in Prism based on the normality of the samples.

For confocal and FLIM imaging of YXCTK2 and importin α-CyPet, cycled egg extracts were prepared, incubated and squashed as described above in which a 20x stock of proteins was added to 1x for final concentrations of 1 μM importin α-CyPet + 150 nM XCTK2; 1 μM importin α-CyPet + 150 nM YPet-XCTK2; 1 μM importin α-CyPet + 100 nM YPet-XCTK2-NLS2b in CSF-XB (10 mM HEPES pH 7.7, 1 mM MgCl_2_, 100 mM KCl, 5 mM EGTA, 50 mM sucrose). Confocal images taken as for Rango-2 except that the X-Rhodamine channel was imaged with 100% gain and the YPet channel at 20% gain. FLIM images were acquired the same as for Rango-2. The overall importin α-CyPet decay data was binned 2×2 and fit to a 3-exponential reconvolution model. The lifetimes per pixel in each image were calculated based on the fitted parameters in which the background and lifetimes were fixed. Images were scaled in SymPhoTime and processed in ImageJ, exported into Excel and graphed in Prism for line scans as described above for Rango-2. The differences between the chromatin and pole lifetimes were calculated in Excel by taking the difference between the average of the three pole lifetimes centered on the peak YXCTK2 and YXNLS fluorescence at both ends of the spindle (positions 3, 4, 5 and 21, 22, 23) from the average of the center three lifetimes (positions 12, 13, 14) for each spindle for each condition and plotted in Prism. ANOVA tests were performed in Prism based on the normality of the samples.

### Reconstitution of XCTK2 interaction with importin α/β and Ran-regulation by FRET

XCTK2 interaction with importin α/β was reconstituted by Förster resonance energy transfer (FRET) using CyPet-XCTK2, importin α-YPet, and importin β. A spectral scan in a Synergy H1 (Bio-Tek) was performed in a Costar #3964 half-area black plate with 100 nM CyPet-XCTK2, 100 nM CyPet-XCTK2 and 200 nM importin α-YPet, or 100 nM CyPet-XCTK2 and 200 nM importin α-YPet with 1.6 μM importin β in 20 mM PIPES pH 6.8, 20 mM KCl, 1 mM MgCl_2_, 1 mM EGTA, 0.1 mM EDTA, 1 mM DTT, 0.2 mg/ml casein. CyPet-XCTK2 was excited at 405 nm and emission recorded from 440-600 nm with 5 nm steps, and a gain of 75. Spectra were background subtracted for YPet fluorescence bleed through and then normalized to the CyPet emission at 460 nm in Excel and plotted in Prism.

XCTK2 Ran-regulation of importin α/β binding was demonstrated by FRET using purified YXCTK2 or YXNLS with purified importin α-CyPet and importin β with and without the addition of purified RanQ69L. A solution of 240 nM YXCTK2 or YXNLS with or without 240 nM importin α-CyPet, 960 nM importin β, 20 mM PIPES pH6.8, 22 mM KCl, 1 mM MgCl_2_, 1 mM EGTA, 0.1 mM EDTA, 0.24 mg/ml, 1 mM DTT was incubated for 10 min in a Costar #3964 half area black plate and scanned spectrally to show binding of importin α/β to XCTK2. Ran control buffer or RanQ69L was then added to 10 μM, the reactions incubated for 10 min, and then scanned again with excitation at 405 nm and emission recorded from 440-600 nm with 5 nm steps, and a gain of 75. Final protein concentrations: 200 nM YXCTK2 or YXNLS, 200 nM importin α-CyPet, 800 nM importin β, 10 μM GST-RanQ69L. Spectra were background subtracted for YPet fluorescence bleed through and then normalized to the CyPet emission at 460 nm in Excel and plotted in Prism.

### XCTK2 importin α/β and microtubule affinity assays

For importin α/β affinity, 100 nM CyPet-XCTK2 (monomer concentration) was mixed with equal volumes of 60 nM-36 μM importin α-YPet or 2 nM-5 μM importin α-YPet with 16 nM-40 μM importin β in 20 mM PIPES pH6.8, 125 mM KCl, 1 mM MgCl_2_, 1 mM EGTA, 0.1 mM EDTA, 0.2 mg/ml casein in a Nunc 384 well non-treated black plate (#264556), incubated 10 min, and scanned in a Synergy H1 (BioTek). A spectral scan was performed in which CyPet was excited at 405 nm, and emission was recorded from 440-600 nm in 5 nm steps with a gain of 150. The spectra were background subtracted for YPet fluorescence and normalized to CyPet emission at 460 nm. Maximum FRET attained with 50 nM CyPet-XCTK2 and 2.5 μM importin α-YPet and 20 μM importin β was a value of 5, which was set as 100% CyPet-XCTK2 bound for fitting to the quadratic equation for one-site binding with ligand depletion in Prism. An extra sum-of-squares F test was performed to compare the *K*_d_ and capacity of importin α/β binding.

To determine the affinity of the tail domain for MTs in the presence and absence of different levels of importin α and/or importin β, purified YPet-Tail was used to circumvent the complexity of the motor domain binding to MTs in addition to the tail domain in the full-length protein. Tubulin was polymerized into MTs at 10 μM tubulin with 0.5 mM GMPCPP (Jena Bioscience) in BRB80/DTT (80 mM PIPES pH6.8, 1 mM MgCl2, 1 mM EGTA, 1 mM DTT) for 20 min at 37°C, paclitaxel added to 10 μM and incubated an additional 10 min. The MTs were then sedimented at 45,000 rpm in a TLA100 rotor (Beckman) for 10 min at 35°C and resuspended in BRB80/DTT, 10 μM paclitaxel at room temperature. A solution of 1 µM YPet-Tail with or without 4 μM or 16 μM importin α/β in 20 mM PIPES pH 6.8, 100 mM KCl, 1 mM MgCl_2_, 1 mM EGTA, 0.1 mM EDTA, 0.4 mg/ml casein was mixed with an equal volume of 0-20 μM MTs in BRB80/DTT, 10 μM paclitaxel to start the reaction. Reactions were incubated for 15 min and sedimented at 45,000 rpm in TLA100 in Optima-TLX at 22°C for 10 min. Supernatants were removed, and the pellet resuspended in an equal volume of resuspension buffer (50 mM PIPES pH6.8, 50 mM, 1 mM MgCl_2_, 1 mM EGTA, 0.05 mM EDTA, 1 mM DTT, 0.2 mg/ml casein, 5 μM Taxol. Equal volumes (15 μl) of the supernatants and pellets were moved to a Nunc 384 well non-treated black plate, and the YPet fluorescence measured in Synergy HI plate reader with 500 nm excitation and emission measured at 530 nm with a gain of 100. The fraction bound (Fract) was determined from amount of YPet fluorescence in the pellet divided by the sum of the supernatant and pellet fluorescence and then normalized to the concentration bound (Fract*0.5 μM), plotted in Prism and fit to the quadratic equation for one-site binding with ligand depletion in Prism. An extra sum-of-squares F test was performed to compare the *K*_d_ and capacity of MT binding.

### Visualization of XCTK2 microtubule cross-linking and sliding by TIRF

To visually characterize XCTK2 parallel and anti-parallel MT cross-linking and sliding, segmented red/green template MTs and green/red cargo MTs were generated by MT extension from GMPCPP MT seeds. Seeds were polymerized from 2 μM tubulin, 1 mM GMPCPP mixes for 30 min and then added 1:10 to 0.5 μM tubulin, 0.5 mM GMPCPP extension mix for 30 min before being sedimented and resuspended in BRB80, 1 mM DTT, 10 μM Taxol. The red/green template MTs were generated from 12% X-rhodamine-tubulin, 10% biotin-tubulin GMPCPP seeds with 10% Alexa488- or Dylight488-tubulin, 10% biotin-tubulin extensions. The green/red cargo MTs were generated from 1% Alexa488- or Dylight488-tubulin GMPCPP seeds with 23% X-rhodamine-tubulin extensions. MT concentration was determined in terms of tubulin dimer concentration by absorbance using the extinction coefficient for tubulin (115,000 M^-1^ cm^-1^).

To visualize sliding, flow chambers were made from biotin-PEG coated #1.5 coverslips and double stick tape as previously described (Ems-McClung et al., 2013). For each reaction condition, a chamber was rinsed with BRB80, 1 mM DTT and then incubated in 5% Pluronic F-127 in BRB80 for 3 min. The chamber was then rinsed with Block (120 mM KCl, 1 μM MgATP, 20 mM PIPES pH6.8, 1 mM MgCl_2_, 1 mM EGTA, 0.1 mM EDTA, 1 mg/ml casein, 10 μM Taxol, 1 mM DTT) and incubated with 0.05 mg/ml NeutrAvidin diluted in Block for 3 min and then rinsed with Block. Template MTs in Block (0.1 μM) were introduced and incubated for 3 min, and then rinsed with Block. YPet-XCTK2 (10 nM) +/-importin α/β (40-80 nM) in 120 mM KCl, 20 mM PIPES pH6.8, 1 mM MgCl2, 1 mM EGTA, 0.1 mM EDTA, 1 mg/ml casein, 5 μM Taxol was flowed in and incubated for 3 min. The chamber was rinsed with Block and then incubated with 0.1 μM cargo MTs in Block for 5 min. To activate sliding prior to imaging, the chamber was then washed with Block containing 5 mM MgATP, 0.32 mg/ml glucose oxidase, 0.055 mg/ml catalase, 25 mM sucrose. Prepared chambers were imaged on a Nikon A1 microscope equipped with a Hamamatsu ORCA Flash 4.0 camera (Melville, NY) controlled by Nikon Elements and an Apo TIRF 100x oil DIC N2 NA 1.49 (WD=120 um) objective for 10 min. The 488 nm argon laser was set to 30% power and the sapphire 561 nm laser set to 15% power. Movies were taken at 15 sec intervals (41 timepoints) as a 2 × 2 grid using the Large Image (λ) setting in Elements with both the red and green channels being acquired for 150 ms before moving to the adjacent field. One to two movies were taken per chamber. Alternatively, movies were imaged on a Nikon 90i microscope with a Nikon Plan Fluor 100x oil 1.3 NA objective and captured with a Photometrics CoolSNAP HQ CCD camera (Roper Scientific, Trenton, NJ) controlled by Molecular Devices MetaMorph (Sunnyvale, CA). Four individual movies were acquired as a multidimensional time-lapse image for 5 min with 30 sec intervals for 500 ms with FITC and TxRed filter sets.

Each Large Image (λ) movie acquired on the Nikon A1 was split into four separate movies prior to sliding analysis. MT cross-linking, sliding, and velocity were scored by hand in Fiji (LOCI, University of Wisconsin-Madison). MT cross-links were scored for orientation (parallel, anti-parallel, or not determined) and type of cross-link (sliding, static, or pivoting). Orientation was determined by the lengths of extensions under the assumption that the plus ends had longer extensions than the minus ends due the differences in dynamic instability for plus and minus ends of MTs (Kristofferson et al., 1986). A MT cross-link was scored as one that slid based on the cargo MTs sliding in reference to a coverslip bound template MT that moved in one direction for least two consecutive frames in the movie. Many MTs moved both directions relative to the template MT. Static cross-linked MTs were identified based on their higher fluorescence intensities relative to nearby template MTs and the presence of a green fluorescent MT seed with red fluorescent MT extensions. Pivoting cross-links were MTs swiveling or waving in and out of the focal plane of the movie, typically at the ends of the MTs. Some MT cross-links displayed multiple types of movement, e.g. sliding to the MT end and then pivoting, or pivoting and then cross-linking along the length of the template MT, and either sliding in different directions or speeds or remaining static. The first sliding or pivoting event observed in the movie was used to characterize the type of cross-link; whereas static cross-links were static throughout the duration of the movie. For each experiment, similar numbers of template MTs (difference < 30) were scored for each condition. The types of MT cross-links were tabulated in Excel as the sum of MTs from two to four separate movies per condition per experiment from six independent experiments and plotted in Prism. Two-tailed unpaired Student’s *t*-tests assuming equal variance or two-tailed Welch’s *t*-tests assuming unequal variance were performed in Prism.

The velocity of MT sliding was measured from the two-color composite movies acquired on the Nikon A1 in Fiji using the MTrackJ plugin. Each sliding and static template-cargo MT cross-link was tracked for the entire length of the movie (10 min) as long as the cargo MT remained cross-linked. Three template MTs in each movie were also tracked to register each movie due to the stage not returning precisely to the same position after acquisition of the four movies per time interval. The tracks were then measured and exported into Excel. To register the measurements of each movie, the average change in x and y for each time interval was determined from the three template MT tracks and then subtracted from x and y position in each cross-link time interval in Excel. The distance the MT moved was then calculated using the Pythagorean theorem. The distances for each movie were plotted, the sliding events identified as described above, and the velocities calculated by dividing by the time interval that the MT slid. Some MT cross-links had multiple sliding events, which were measured and included as individual sliding events in the velocity results. Velocities for each orientation were then tabulated per condition and graphed as a frequency histogram based on the number of values with the best fit log Gaussian curve. Statistical difference was determined using the extra sum-of-squares F-test for the geometric mean of the log Gaussian distribution of velocities. The number of sliding events per MT cross-link was determined per condition, tabulated, and graphed in Prism with the mean and SD indicated. The sliding events per MT cross-link were compared using the two-tailed Welch’s unpaired *t*-test in Prism.

## Online supplemental materials

Fig. S1 shows the affinity of XCTK2 CXCTK2 to importin α and importin α/β as well as the MT affinity of the XCTK2 tail domain in the presence and absence of excess of importin α/β. Fig. S2 shows the morphology of the spindles with enhanced RanGTP gradients that were stained with α-XCTK2, imaged, and analyzed in Fig. 4 D-H as well as the localization of NuMA in enhanced RanGTP gradients that were imaged and analyzed similar to those stained for XCTK2 localization in Fig. 4. Table S1-4 document the means and SD or SEM for each analysis in Figs. 2-4, the statistical analysis used, the number of samples, and the number of independent experiments. Videos 1 and 2 show the *in vitro* cross-linking and sliding assay performed in Fig. 3.

**Figure S1.**
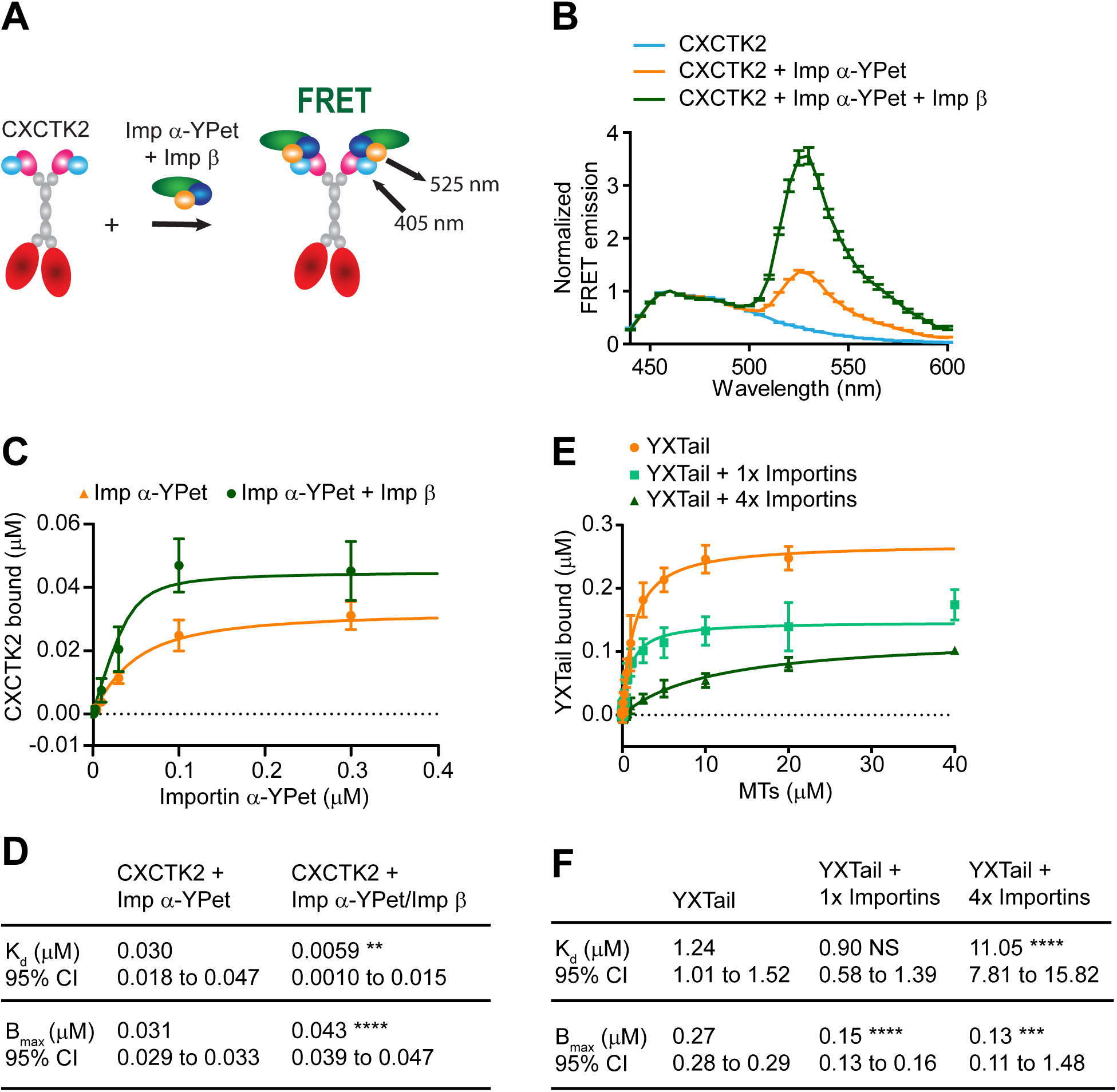
XCTK2 tail MT binding is tunable by importin α/β association. **(A)** Schematic of FRET biosensors to detect XCTK2 and importin α/β association where XCTK2 is N-terminally tagged with CyPet (CXCTK2) and importin α is C-terminally tagged with YPet (imp α-YPet). **(B)** Solution-based FRET assay of CXCTK2 with importin α-YPet ± importin β. The normalized FRET ratios are graphed as mean ± SEM. from 440-600 nm (*n* = 4-5 independent experiments). **(C**, **D)** Affinity assays of CXCTK2 with importin α-YPet ± importin β. The mean ± SD of CXCTK2 bound for each importin α-YPet concentration and the best-fit quadratic binding curves are graphed from **(C)** with the binding data summarized in **(D)** (*n* = 4-5 independent experiments; extra sum-of-squares F-test compared to CXCTK2 + imp α-YPet: *K*_d_, *P* = 0.0088; B_max_, *P* < 0.0001). CXCTK2 bound is graphed from 0-0.3 μM Imp α-YPet, excluding 1 and 2.5 μM Imp α-YPet data points. **(E**, **F)** MT affinity assays of YXTail in the absence and presence of equal molar (1x) or four-fold excess (4×) of importin α/β. The amount of YXTail bound is plotted as the mean ± SD for each MT concentration, and the best-fit quadratic binding curves are graphed **(E)** with the binding data summarized in **F** (*n* = 5-8 independent experiments; extra sum-of-squares F-test compared to YXTail alone: YXTail + 1x imp α/β, *K*_d_ NS, not significant, B_max_ *P* < 0.0001; YXTail + 4× imp α/β, *K*_d_ *P* < 0.0001, *P* = 0.0002).

**Figure S2.**
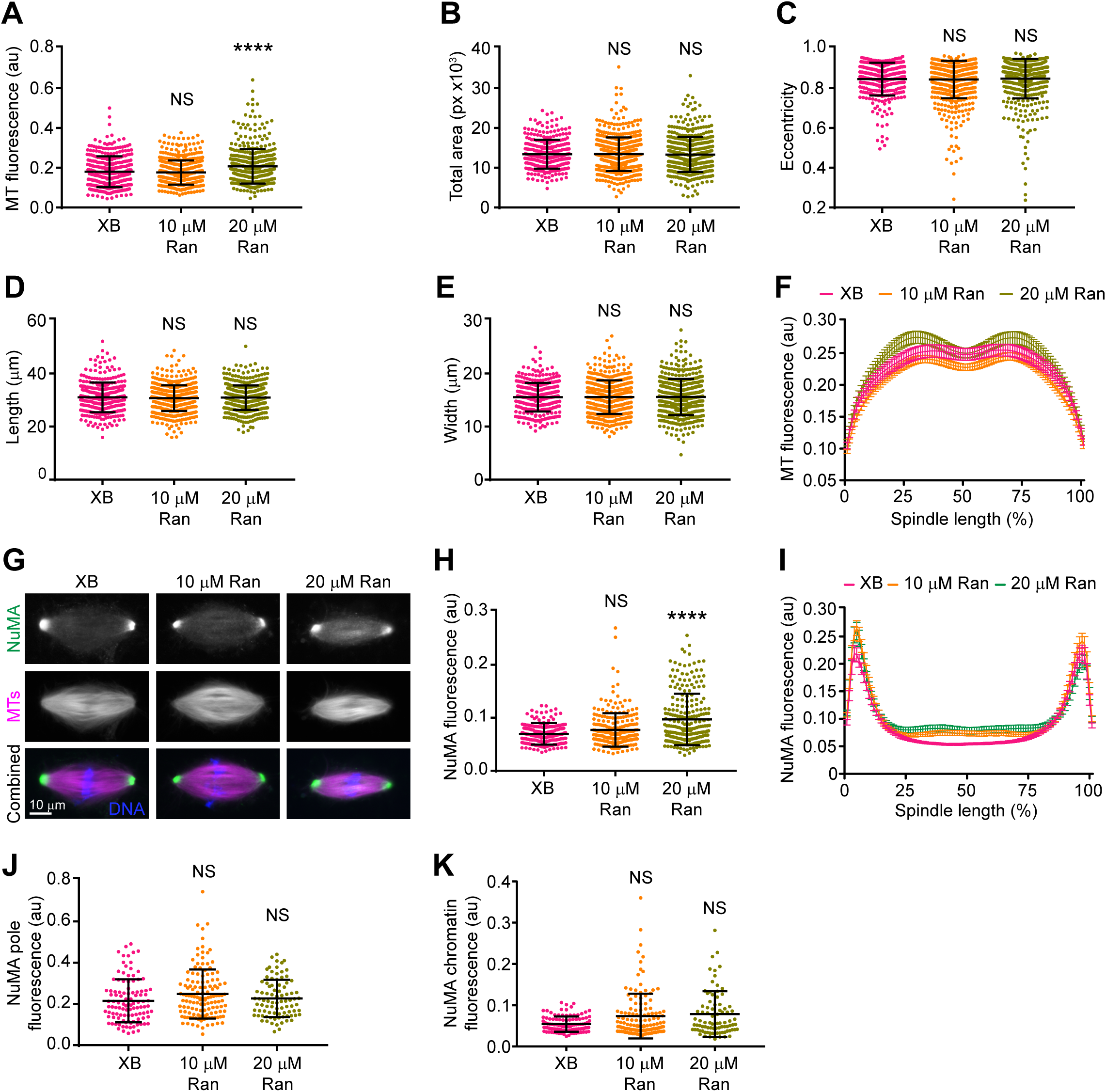
Enhancing the RanGTP gradient does not change spindle morphology or NuMA localization. **(A-E)** Spindle morphology analysis with Cell Profiler of spindles measured in **Fig. 4 D** (*n* = 314-564 spindles per condition from 4-5 independent experiments; Kruskal-Wallis test with Dunnett’s Multiple Comparisons test compared to XB control buffer, NS, not significant, **** *P* < 0.0001). **(F)** Line scan analysis of MT polymer distribution of spindles analyzed in **Fig. 4 F** graphed as the mean ± SEM. **(G)** Representative wide-field fluorescence microscopy images of spindle assembly reactions with XB control buffer, 10 μM Ran, or 20 μM Ran addition stained with α-NuMA that were performed in parallel to those in **Fig. 4**. Scale bar: 10 μm. **(H)** Total α-NuMA fluorescence of spindles assembled in **G** and plotted with the mean ± SD (*n* = 209-240 spindles per condition from 4 independent experiments; Kruskal-Wallis test with Dunnett’s Multiple Comparisons test compared to XB control buffer, NS, not significant, \*\*\*\**P* < 0.0001). **(I)** Line scan analysis of bipolar spindles analyzed in **G**, normalized for percent spindle length (101 bins), and graphed as the mean ± SEM (*n* = 86-134 spindles per condition from 4 independent experiments). **J** and **K**, NuMA peak pole fluorescence and chromatin fluorescence as calculated in **Fig. 4, G** and **H** and plotted with the mean ± SD indicated (Kruskal-Wallis test with Dunnett’s Multiple Comparisons test compared to XB buffer control, NS, not significant).

Video 1. **YXCTK2 slides both anti-parallel (left) and parallel (right) MTs**. *In vitro* time-lapse TIRF microscopy assay of 10 nM YXCTK2 cross-linking and sliding dual color segmented MTs shown in Fig. 3 B. Images were collected for 10 min at 15 sec intervals for 41 frames and are shown at 7 frames per sec. Template MTs have magenta (X-Rhodamine) MT seeds with green (DyLight488) MT extensions and cargo MTs have green MT seeds and magenta MT extensions. Template MT plus (+) and minus (-) ends are labeled, and the cargo MT plus end is labeled with an asterisk.

Video 2. **Cargo MTs cross-linked with YXCTK2 in the presence of importin α/β have more frequent sliding events**. *In vitro* time-lapse TIRF microscopy assay of 10 nM YXCTK2 + 80 nM importin α/β cross-linking and sliding dual color segmented MTs shown in Fig. 3 B. Images were collected as for Video 1 for anti-parallel (left) and parallel (right) MT cross-links. Template MT plus (+) and minus (-) ends are labeled, and the cargo MT plus end is labeled with an asterisk.

**Table S1.** Importin α-CyPet lifetime chromatin to pole differences.

| | Importin<br>α-CyPet | YXNLS | YXCTK2 |
| --- | --- | --- | --- |
| **Importin α-CyPet Chromatin to Pole Lifetime Difference (ns)** | -0.02905 | -0.02383 | 0.02289 |
| SD | 0.02903 | 0.03273 | 0.04838 |
| One-way ANOVA with Tukey's multiple comparisons test |  | 0.8537 | <0.0001 |
| Spindle Number | 38 | 30 | 58 |
| Experiment number | 3 | 3 | 3 |

Mean chromatin to pole lifetime differences measured from the line scans in Fig. 2 C and graphed in Fig. 2 E.

**Table S2.** YXCTK2 ± importin α/β MT cross-linking and sliding.

|  | YXCTK2 | YXCTK2 +<br>Importin α/β | P value |
| --- | --- | --- | --- |
| **Template MTs** | 442 | 451 | 0.9403 |
| SD | 189 | 194 |  |
| Total number | 2654 | 2705 |  |
| **Cross-linked MTs** | 122 | 59 | 0.0109 |
| SD | 44 | 23 |  |
| Total number | 729 | 352 |  |
| **Sliding cross-linked MTs** | 78 | 31 | 0.0230 |
| SD | 35 | 26 |  |
| Total number | 470 | 184 |  |
| **Static cross-linked MTs** | 24 | 19 | 0.4709 |
| SD | 13 | 10 |  |
| Total number | 144 | 112 |  |
| **Pivoting cross-linked MTs** | 19 | 10 | 0.0373 |
| SD | 9 | 4 |  |
| Total number | 116 | 58 |  |
| **Anti-parallel cross-linked MTs** | 11 | 5 | 0.0437 |
| SD | 5 | 3 |  |
| Total number | 63 | 31 |  |
| **Parallel cross-linked MTs** | 14 | 9 | 0.1705 |
| SD | 6 | 4 |  |
| Total number | 81 | 56 |  |
| **Anti-parallel sliding cross-linked MTs** | 8 | 3 | 0.0620 |
| SD | 5 | 2 |  |
| Total number | 48 | 19 |  |
| **Anti-parallel static cross-linked MTs** | 3 | 2 | 0.6463 |
| SD | 2 | 1 |  |
| Total number | 15 | 12 |  |
| **Parallel sliding cross-linked MTs** | 12 | 6 | 0.0899 |
| SD | 7 | 3 |  |
| Total number | 69 | 36 |  |
| **Parallel static cross-linked MTs** | 2 | 3 | 0.4075 |
| SD | 2 | 3 |  |
| Total number | 12 | 20 |  |
| Experiment number | 6 | 6 |  |
| **Anti-parallel velocity of sliding (nm/s)** | 14.55 | 12.90 | 0.4299 |
| Geometric SD | 2.71 | 1.89 |  |
| Total Sliding Events | 42 | 60 |  |
| **Parallel velocity of sliding (nm/s)** | 8.55 | 9.88 | 0.4020 |
| Geometric SD | 3.06 | 2.14 |  |
| Total Sliding Events | 71 | 116 |  |
| Experiment number | 3 | 3 |  |
| **Sliding events per MT cross-link** | 2.40 | 5.13 | <0.0001 |
| SD | 1.77 | 2.97 |  |
| Total MT cross-links | 212 | 140 |  |
| Experiment number | 3 | 3 |  |

**Table S3.** Rango-2 lifetime pole to chromatin differences.

|  | **XB** | **10 μM Ran** | **20 μM Ran** |
| --- | --- | --- | --- |
| **Rango-2 pole to chromatin lifetime difference (ns)** | 0.04247 | 0.05184 | 0.05115 |
| SD | 0.02217 | 0.01879 | 0.02317 |
| One-way ANOVA with Tukey's multiple comparisons test |  | 0.0372 | 0.0351 |
| Spindle number | 78 | 57 | 78 |
| Experiment number | 4 | 3 | 4 |

**Table S4.** Fluorescence and morphology of spindles assembled in enhanced RanGTP gradients in *Xenopus* egg extracts.

|  | **XB** | **10 μM Ran** | **20 μM Ran** |
| --- | --- | --- | --- |
| **XCTK2 fluorescence (au)** | 0.1205 | 0.1369 | 0.1451 |
| SD | 0.03693 | 0.03809 | 0.04572 |
| Kruskal-Wallis with Dunn's multiple comparisons test |  | <0.0001 | <0.0001 |
| Spindle number | 314 | 564 | 487 |
| Experiment number | 5 | 5 | 5 |
| **MT fluorescence (au)** | 0.1801 | 0.1763 | 0.2069 |
| SD | 0.07748 | 0.06103 | 0.08669 |
| Kruskal-Wallis with Dunn's multiple comparisons test |  | >0.9999 | <0.0001 |
| Spindle number | 328 | 509 | 453 |
| Experiment number | 4 | 4 | 4 |
| **Total area (px)** | 12697 | 12935 | 12540 |
| SD | 3278 | 3924 | 3781 |
| Kruskal-Wallis with Dunn's multiple comparisons test |  | >0.9999 | >0.9999 |
| **Eccentricity** | 0.8712 | 0.8755 | 0.8876 |
| SD | 0.07026 | 0.05873 | 0.04387 |
| Kruskal-Wallis with Dunn's multiple comparisons test |  | >0.9999 | 0.4134 |
| **Length (μm)** | 30.45 | 30.64 | 30.41 |
| SD | 5.368 | 4.562 | 4.331 |
| Kruskal-Wallis with Dunn's multiple comparisons test |  | >0.9999 | >0.9999 |
| **Width (μm)** | 15.63 | 15.21 | 15.34 |
| SD | 2.854 | 3.617 | 3.423 |
| Kruskal-Wallis with Dunn's multiple comparisons test |  | >0.9999 | >0.9999 |
| Spindle number | 314 | 564 | 487 |
| Experiment number | 5 | 5 | 5 |
| **XCTK2 pole fluorescence (au)** | 0.1569 | 0.1862 | 0.1998 |
| SD | 0.04905 | 0.05667 | 0.06651 |
| Kruskal-Wallis with Dunn's multiple comparisons test |  | <0.0001 | <0.0001 |
| **XCTK2 chromatin fluorescence (au)** | 0.1262 | 0.1421 | 0.1521 |
| SD | 0.03553 | 0.04858 | 0.05817 |
| Kruskal-Wallis with Dunn's multiple comparisons test |  | 0.0424 | <0.0001 |
| Spindle number | 168 | 243 | 248 |
| Experiment number | 4 | 4 | 4 |
| **NuMA fluorescence (au)** | 0.06903 | 0.07720 | 0.09696 |
| SD | 0.01803 | 0.03147 | 0.04846 |
| Kruskal-Wallis with Dunn's multiple comparisons test |  | 0.1527 | <0.0001 |
| Spindle number | 209 | 240 | 239 |
| Experiment number | 4 | 4 | 4 |
| **NuMA pole fluorescence (au)** | 0.2153 | 0.2259 | 0.2269 |
| SD | 0.1045 | 0.09043 | 0.09038 |
| Kruskal-Wallis with Dunn's multiple comparisons test |  | >0.9999 | >0.9999 |
| **NuMA chromatin fluorescence (au)** | 0.05438 | 0.07358 | 0.07644 |
| SD | 0.01853 | 0.05421 | 0.05276 |
| Kruskal-Wallis with Dunn's Multiple Comparisons test |  | >0.9999 | >0.9999 |
| Spindle number | 107 | 134 | 86 |
| Experiment number | 4 | 4 | 4 |

## Acknowledgements

The authors would like to thank Ben Walker, Kelly Hartsough, and Rebecca Heald for critical comments on the manuscript, and all members of the Walczak lab for helpful discussion. We thank Petr Kaláb and Sid Shaw for help on the FLIM analysis and Jim Powers for his help with multiple types of imaging. This work was supported by NIH grants GM059618 and GM122482 to CEW and GM099309 to LNW. The Light Microscopy Imaging Center is supported in part by the OVPR, COAS, IUSM-BL, and the School of Optometry at Indiana University.

## Author contributions

S.C.E designed the study, performed most of the experiments, and analyzed the data. S.Z. performed the immunofluorescence assays of spindle assembly experiments with Ran addition. M.E., S.Z, and S.M. contributed to data collection and image analysis. L.N.W. contributed reagents and design advice to the study. C.E.W contributed to the design and interpretation of the study, as well as writing and editing of the manuscript.

